# Improved correspondence of fMRI visual field localizer data after cortex-based macroanatomical alignment

**DOI:** 10.1101/2021.02.26.433066

**Authors:** Mishal Qubad, Catherine V. Barnes-Scheufler, Michael Schaum, Eva Raspor, Lara Rösler, Benjamin Peters, Carmen Schiweck, Rainer Goebel, Andreas Reif, Robert A. Bittner

## Abstract

Studying the visual system with fMRI often requires using localizer paradigms to define regions of interest (ROIs). However, the considerable interindividual variability of the cerebral cortex represents a crucial confound for group-level analyses. Cortex-based alignment (CBA) techniques reliably reduce interindividual macroanatomical variability. Yet, their utility has not been assessed for visual field localizer paradigms, which map specific parts of the visual field within retinotopically organized visual areas. We evaluated CBA for an attention-enhanced visual field localizer mapping homologous parts of each visual quadrant in 50 participants. We compared CBA with volume-based alignment and a surface-based analysis, which did not include macroanatomical alignment. CBA led to the strongest increase in the probability of activation overlap (up to 40 percent). On the group level, CBA led to the most consistent increase in ROI size while preserving vertical ROI symmetry. Overall, our results indicate, that in addition to the increased signal-to-noise ratio of a surface-based analysis macroanatomical alignment considerably improves statistical power. These findings confirm and extend the utility of CBA for the study of the visual system in the context of group analyses. CBA should be particularly relevant when studying neuropsychiatric disorders with abnormally increased interindividual macroanatomical variability.

## INTRODUCTION

The visual system includes a multitude of topographical representations of varying resolution across increasingly specialized visual areas^1^. Functional magnetic resonance imaging (fMRI) offers a variety of methods either to map these topographical representations in full, or to localize specific visual areas or retinotopic positions within their topography. These approaches are essential not only for the fine-grained study of fundamental properties of the visual system^1^, but also for investigating the role of these areas for higher-order cognitive processes such as visual attention and working memory^2–6^. This also extends to translational studies of visual dysfunction and its cognitive consequences in neuropsychiatric disorders^7, 8^.

Methods for fMRI-based visual mapping, i.e., techniques to define regions of interest in the visual system based on specific functional properties, fall in in three broad categories: retinotopic mapping, visual field localizer and functional localizer paradigms. Retinotopic mapping and the more advanced population receptive field mapping allow the complete delineation of early visual areas^1, 9, 10^. Conversely, visual field localizer paradigms can map a circumscribed region within a retinotopically organized visual area^11, 12^. Finally, functional localizers can detect higher-order visual areas such as the fusiform face area, parahippocampal place area (PPA), extrastriate body area (EBA) and lateral occipital complex (LOC), which are clustered and show specialization for the processing of specific categories of complex visual information^1, 13, 14^. In most fMRI studies, high interindividual anatomical variability of cortical areas in terms of both size and location constitutes an important challenge^15–23^. For instance, it has been shown that primary visual cortex (V1) can differ in size by about 2-fold between individuals^17^. Furthermore, anatomical variability in terms of location has been shown to be particularly pronounced in extra striate visual areas^24^. This crucial confound reduces the power to reliably map visual areas at the group level.

One way to mitigate this problem for the visual system is to pool single-subject regions of interest (ROIs), while simultaneously using the overall group-based probability for that ROI at each point in a Cartesian coordinate system as a constraint^25–27^. While such a single-subject-based analysis improves sensitivity and functional resolution compared to a standard group-based approach, it does not actually reduce macroanatomical variability. Additionally, studying the interplay between visual areas and other cortical areas more directly involved in higher-order cognitive processes with whole-brain methods such as functional connectomics network analyses^28^ might preclude a single-subject based strategy.

Such group-based analyses typically require spatial normalization of structural and functional imaging data to a common Cartesian coordinate system such as Talairach^29^ or MNI^30^ space. In its most basic form, volume-based spatial normalization employs a linear transformation that matches the overall extent of the brains to a standard brain template. While transformation into Talairach space relies on anatomical landmarks, transformation into MNI space utilizes fully data driven registration of structural images to an average template brain^30^. While these spatial normalization approaches inherently lead to an alignment of brains, the underlying algorithms are not geared specifically toward aligning homologous brain structures. Conversely, more refined methods employ non-linear warping algorithms guided by intensity differences to improve macroanatomical alignment^31^. Thus, all of these methods can be categorized as volume-based alignment (VBA) techniques. However, both linear and nonlinear VBA mostly disregard the topological properties of the cerebral cortex and its geometric features such as sulci and gyri. Consequently, VBA methods result in a considerable amount of residual interindividual anatomical variability^32, 33^. Surface-based procedures constitute an important alternative approach. Surface-based spatial normalization typically uses a geodesic coordinate system, which allows for a two-dimensional representation of the cerebral cortex and respects the cortical topography to a much larger degree than traditional Cartesian coordinate systems^18, 34^. This approach offers two main advantages over VBA. First, surface-based spatial normalization allows to constrain data readout and data preprocessing such as spatial smoothing to cortical tissue, greatly reducing signal contamination by white matter and cerebrospinal fluid. It also avoids contamination from cortical areas proximal in volume space but considerably more distant in surface space. Consequently, spatial smoothing in surface space is superior to spatial smoothing in volume space^19, 35, 36^. The second advantage of surface-based spatial normalization is the possibility to use individual cortical folding patterns for an additional, fully data-driven macroanatomical alignment of the cerebral cortex^34^. Compared to VBA techniques, these cortex-based alignment (CBA) methods considerably improve anatomical correspondence of cortical structures while respecting cytoarchitectonic boundaries^37^. Thus, CBA leads to a notable reduction of inter-individual anatomical variability^18, 34, 38–40^.

Importantly, previous studies have often exclusively compared surface-based data before and after macroanatomical alignment^19, 41^, essentially using the former approach as a proxy for VBA. Yet, this comparison only reflects the second advantage of CBA, namely the use of macroanatomical alignment instead of VBA. However, in this case both data sets benefit equally from reduced signal contamination, likely underestimating the full effects of CBA. Assessing the impact of this first advantage of surface-based analyses in isolation requires a comparison of VBA with a surface-based analysis without macroanatomical alignment. We refer to this intermediate approach as a “surface-based analysis using VBA” (SBAV). Thus, studying both advantages of CBA requires the comparison of three approaches: VBA, SBAV and CBA.

Due to the advantageous properties outlined above, CBA methods have been proposed as an alternative approach to VBA specifically for the visual system^26^. Several studies have compared the impact of VBA and CBA methods on specific visual mapping techniques. For retinotopic mapping an improvement of functional overlap in both V1 and V2 after CBA has been demonstrated^34, 42^. Furthermore, for functional localizer data CBA substantially increases the overlap of object processing areas LOC, FFA and PPA across subjects^19, 43–45^. Conversely, the effects of CBA on visual field localizer paradigms mapping specific retinotopic positions have not been studied. Thus, the utility of CBA has been demonstrated for two of the three main categories of visual mapping methods, i.e., those methods, which map whole areas, either defined by cytoarchitectonic (e.g. V1) or functional (e.g. FFA) properties. Conversely, it remains unclear, to which degree CBA can improve the alignment of ROIs mapped by visual field localizer paradigms. Such paradigms are required for the detailed study of the local processing of simple visual stimuli in early visual areas^11, 12, 46–48^. Flashing checkerboards covering the exact area of interest within the visual field are primarily used for this purpose. Checkerboards lead to a particularly strong BOLD-signal increase in early visual areas (V1-V3)^49^. To maximize the fidelity of the resulting localizer maps, visual field localizer paradigms typically utilize the fact that attentional modulation induced by task demands significantly enhances response reliability across visual areas. This can be accomplished by adding a simple target detection task^50^.

We used such an attention-enhanced visual field localizer paradigm to map a circumscribed location in each visual quadrant across early visual areas aiming to define ROIs to be used for the study of higher cognitive processes. We chose a CBA method using a dynamic group average as the target brain^19^, thus eliminating the possible confound of a static CBA target based on an individual brain, whose cortical folding pattern might by chance deviate considerably from the group average. Our primary goal was to examine the effects of CBA for a visual field localizer paradigm. More specifically, we aimed to determine, whether macroanatomical alignment improves the reliability of mapping subregions within retinotopically organized visual areas delineated by such a paradigm at the group level. To this end, in addition to the analysis of the full single-subject ROIs, we also examined the correspondence of single-subject ROI peak vertices, i.e., single vertices showing the strongest level of activation in each subject for each visual quadrant. We conducted this analysis, because peak vertices are a good approximation of the center of a ROI and thus allow for a more precise assessment and visualization of the effects of CBA. Based on previous findings for other localizer paradigm classes and the relatively good structural-functional correspondence in posterior occipital cortex, we expected to observe a benefit of CBA compared to SBAV when aligning subregions within early visual cortex for both full ROIs and peak vertices. Our second goal was to examine the effects of SBAV. More specifically, we aimed to assess the impact of surface-based functional data readout and pre-processing without macroanatomical alignment. Here, we expected a general improvement of the signal-to-noise ratio for SBAV compared to VBA and a corresponding global increase in ROI size for all visual quadrants.

Notably, several studies have shown differential response properties such as receptive field size by visual quadrant or hemifield for homologous early visual areas. For instance, improved behavioral performance and higher BOLD-signal amplitudes were observed in the lower visual hemifield^51–54^. We were therefore also interested, whether we could observe differences between upper and lower visual hemifields in our group analysis after CBA. Overall, the aim of the study was to close an important gap in the evaluation of CBA for the study of the visual system. Since visual field localizers are crucial for investigating contributions of the visual system to higher-order cognitive processes, our results should have implications for the study of visual cognition in both basic and translational research.

## RESULTS

### Visual quadrant ROIs (group level)

Group-level mapping of the four visual quadrants revealed notable differences for the three alignment techniques (VBA, SBAV, CBA) (Figure 4, Table 1). For the lower right visual quadrant, ROI size increased considerably from VBA to SBAV, but decreased for CBA. For the lower left visual quadrant, ROI size decreased slightly from VBA to SBAV, but increased considerably for CBA. For the upper left visual quadrant, ROI size increased considerably from VBA to SBAV and increased further for CBA. For the upper right visual quadrant ROI size increased slightly from VBA to SBAV and increased considerably for CBA. Thus, three out of four visual quadrant ROIs exhibited a pattern of continuously increasing cluster size reflecting an increasing extent of significant position selectivity across alignment techniques. Conversely, the ROI of the lower right visual quadrant showed a cluster size decrease after CBA. For SBAV this ROI also showed by far the greatest extent of any ROI even encompassing posterior parts of temporal cortex.

**Table 1.**
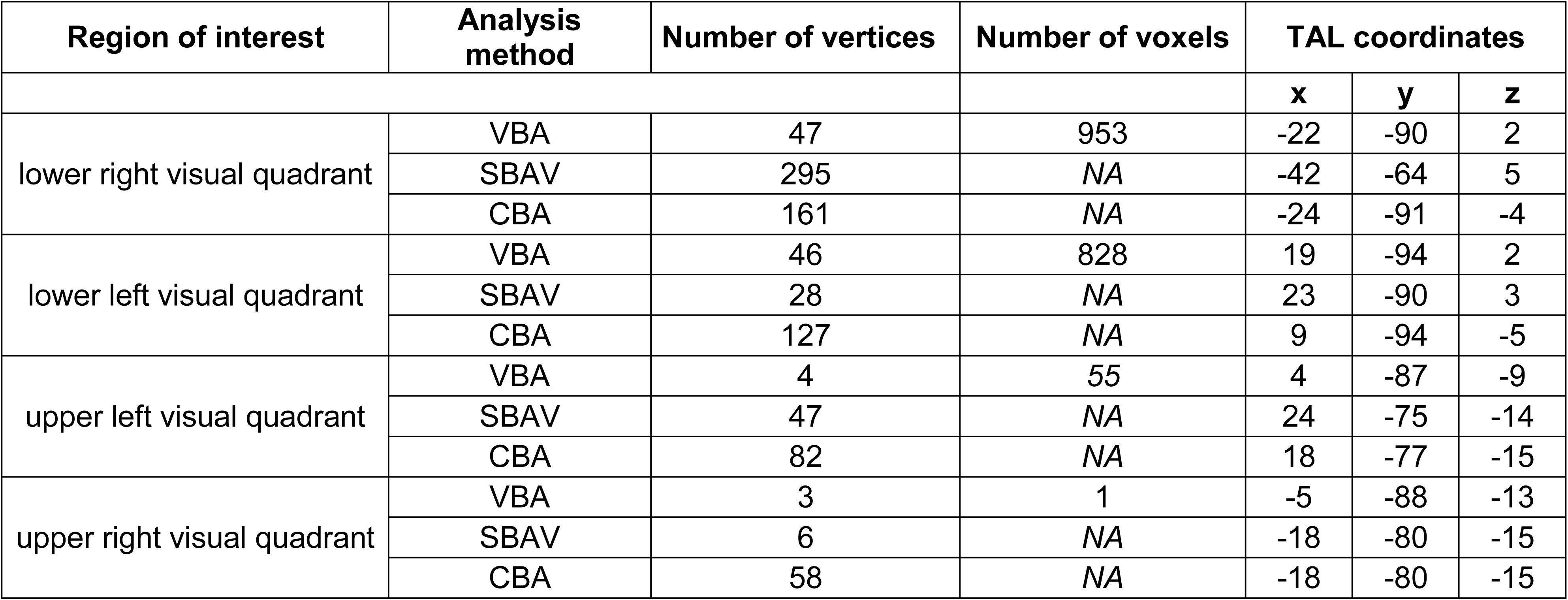
Group ROIs. Size and Talairach coordinates of the group ROIs of the corresponding visual quadrants for the VBA, SBAV and CBA data sets. Three out of four visual quadrant ROIs showed a consistent pattern of increasing cluster, i.e., an increasing extent of position selectivity across alignment techniques.

Within group ROIs, average time courses showed clear position selectivity, which was not affected by alignment technique as indicated by the negative results of our linear mixed models (Table 2). Notably, asymmetry indices (AIs) revealed markedly greater vertical symmetry of both upper and lower hemifield ROIs for VBA and CBA compared to SBAV (Table 3). After CBA, ROI sizes for the lower visual hemifield were considerably larger size than for the upper visual hemifield (Table 1).

**Table 2.** Linear mixed model results. To test for differences in the strength of position selectivity within corresponding ROIs across alignment techniques, we conducted separate linear mixed models with random intercept for each visual quadrant. We used each subject’s t-values as the dependent variable and the alignment techniques (VBA, SBAV and CBA) as the independent variable. We adjusted p-values using Bonferroni correction. We did not observe any significant effect, indicating that this measure off position selectivity was not affected by alignment technique. * Random effect variance estimate at the subject-level for the LR ROI was 0, resulting in a singular fit when using lmer. Therefore, results for LR were estimated without a random intercept for ID and are thus equivalent to a regular ANOVA.

|  | **Sum Sq** | **Mean Sq** | **NumDF** | **DenDf** | **F value** | **Pr (>F)** | **p-value (corr.)** |
| --- | --- | --- | --- | --- | --- | --- | --- |
| **LR** | 183.2 | 92 | 2 | 147 | 0.31 | 0.737 | 1.000 |
| **LL** | 2660.2 | 1330.1 | 2 | 98 | 1.25 | 0.291 | 1.000 |
| **UL** | 2546.2 | 1273.1 | 2 | 98 | 3.36 | 0.039 | 0.156 |
| **UR** | 73678 | 36839 | 2 | 98 | 1.19 | 0.308 | 1.000 |

**Table 3.**
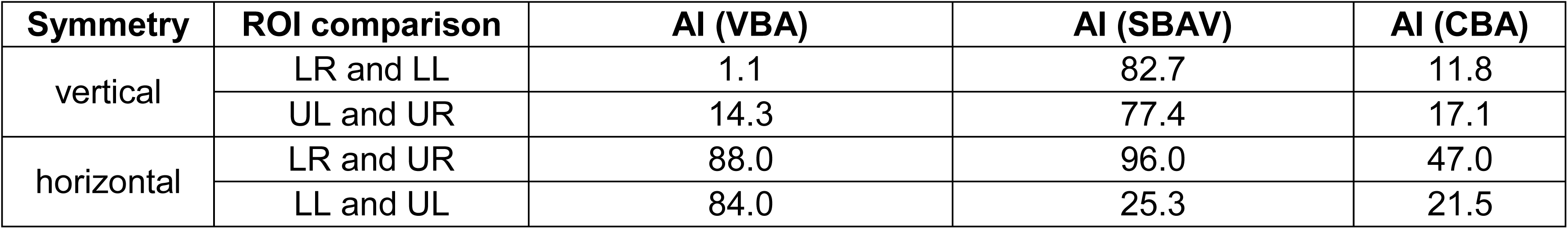
Vertical and horizontal asymmetry indices (AIs). To assess the impact of the three alignment techniques on horizontal and vertical symmetry of our group-level ROIs, we computed an ROI size asymmetry index between each pair of ROIs. AIs revealed greater vertical symmetry of the upper and lower hemifield ROIs for VBA and CBA compared to SBAV.

| **Symmetry** | **ROI comparison** | **AI (VBA)** | **AI (SBAV)** | **AI (CBA)** |
| --- | --- | --- | --- | --- |
| vertical | LR and LL | 1.1 | 82.7 | 11.8 |
|  | UL and UR | 14.3 | 77.4 | 17.1 |
| horizontal | LR and UR | 88.0 | 96.0 | 47.0 |
|  | LL and UL | 84.0 | 25.3 | 21.5 |

### Probability maps

For all three data sets, the maximum probability of activation overlap was consistently located at the center of each ROI as defined in our previous group analysis (Figure 4). For VBA data, probability maps (PMs) showed a relatively wide spread of functional activation around the core ROIs (Figure 5a, Table 4). Maximum probability of activation overlap was approximately 50 percent. For SBAV data, PMs showed an even wider spread of functional activation around the core ROIs (Figure 5b, Table 4). Maximum probability of activation overlap was approximately 45 percent. For CBA data, PMs showed a noticeable decrease in the spread of functional activation around the core ROIs with a corresponding increase in the maximum probability of overlap at the center of the core ROIs (Figure 5c, Table 4). Maximum probability of activation overlap was approximately 75 percent.

**Table 4.**
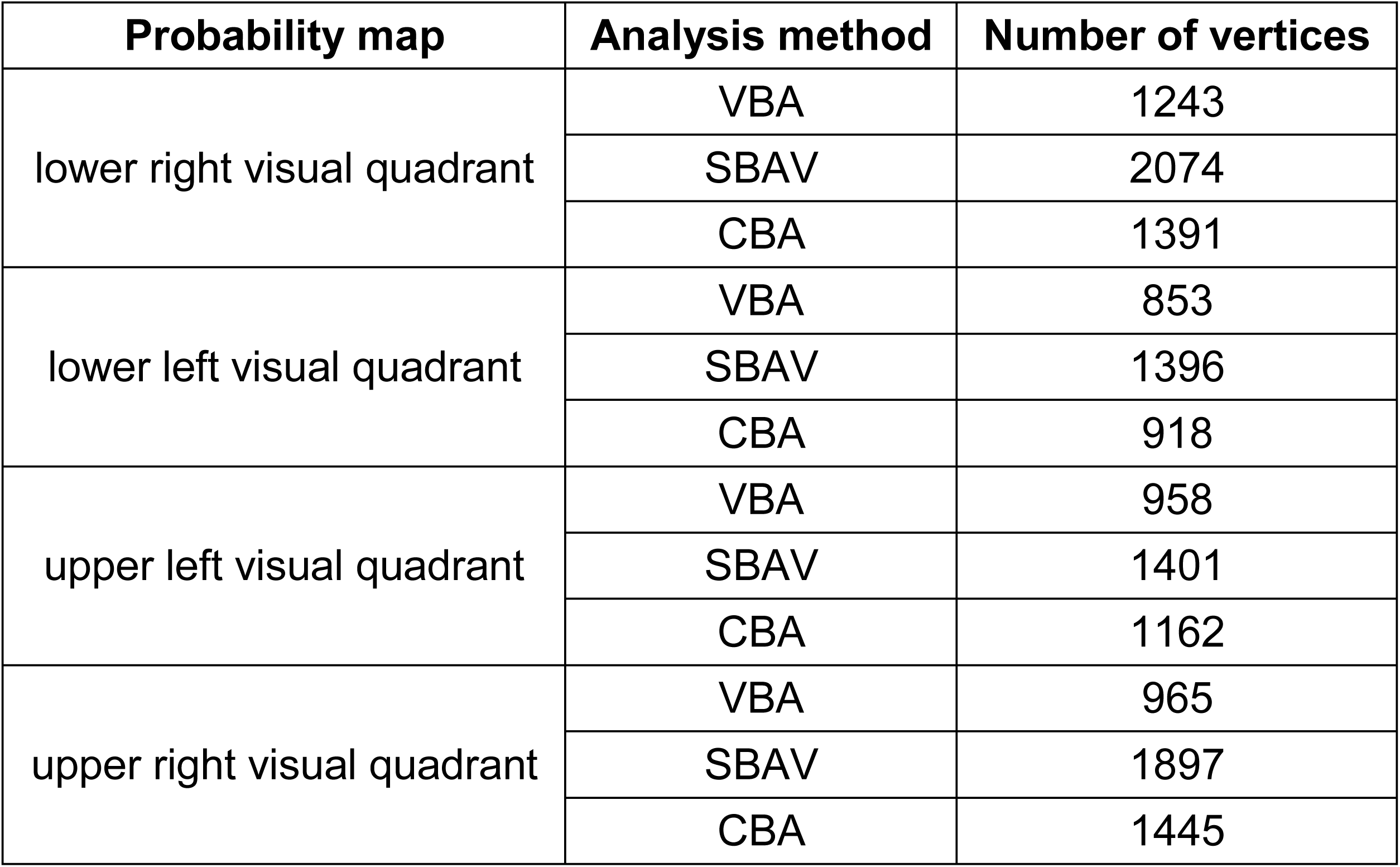
Extent of probability maps. For each visual quadrant and analysis methods, we counted the number of vertices in the corresponding probability maps exceeding the threshold of 10 percent probability of activation overlap. For VBA, maximum probability of activation overlap was around 50 percent. For SBAV, maximum probability of activation overlap was around 45 percent. For CBA, maximum probability of activation overlap was around 75 percent.

| Probability map | Analysis method | Number of vertices |
| --- | --- | --- |
| lower right visual quadrant | VBA | 1243 |
|  | SBAV | 2074 |
|  | CBA | 1391 |
| lower left visual quadrant | VBA | 853 |
|  | SBAV | 1396 |
|  | CBA | 918 |
| upper left visual quadrant | VBA | 958 |
|  | SBAV | 1401 |
|  | CBA | 1162 |
| upper right visual quadrant | VBA | 965 |
|  | SBAV | 1897 |
|  | CBA | 1445 |

### Probability difference maps

Probability difference maps (PDMs) revealed a differential impact of the individual methodological elements of our overall macroanatomical alignment approach. For the impact of surface-based functional data readout and pre-processing compared to standard volume-based alignment, the corresponding PDM (SBAV minus VBA) showed an increase in the probability of activation overlap of up to 15 percent around the central ROIs. Conversely, at the location of the central ROIs we mostly observed a decrease in the probability of activation overlap of up to 10 percent (Figure 6a, Table 5). Notably, changes were widespread, partly extending into posterior temporal and parietal cortex. For the additional impact of macroanatomical alignment, the corresponding PDM (CBA minus SBAV) showed an increase in the probability of activation overlap of up to 35 percent in the central ROIs (Figure 6b, Table 5). Conversely, more peripheral occipital regions showed a decrease in the probability of activation overlap of up to 35 percent. Overall, changes were considerably less wide-spread than for the SBAV minus VBA comparison. For the additive impact of both methodological elements, the corresponding PDM (CBA minus VBA) showed an increase in the probability of activation overlap of up to 30 percent in the central ROIs (Figure 6c, Table 5). Conversely, more peripheral occipital regions as well as posterior temporal and parietal cortex showed a decrease in the probability of activation overlap of up to 20 percent. Overall, the spatial extent of these effects fell in between that of the other two comparisons.

**Table 5.** Extent of probability difference maps (PDMs). For each visual quadrant and comparison between analysis methods (SBAV minus VBA, CBA minus SBAV, CBA minus VBA), we counted the number of vertices in the corresponding PDMs exceeding the threshold of 5 percent difference in probability of activation overlap. Overall, the extent of PDMs was consistently greatest for the CBA minus SBAV comparison, i.e., the isolated effect of effect of macroanatomical alignment.

| Probability difference map | Analysis method | Number of vertices |
| --- | --- | --- |
| lower right visual quadrant | SBAV minus VBA | 1169 |
|  | CBA minus SBAV | 1341 |
|  | CBA minus VBA | 739 |
| lower left visual quadrant | SBAV minus VBA | 570 |
|  | CBA minus SBAV | 694 |
|  | CBA minus VBA | 402 |
| upper left visual quadrant | SBAV minus VBA | 612 |
|  | CBA minus SBAV | 680 |
|  | CBA minus VBA | 546 |
| upper right visual quadrant | SBAV minus VBA | 689 |
|  | CBA minus SBAV | 964 |
|  | CBA minus VBA | 795 |

### Spatial variability of ROI peak vertex distribution (single-subject level)

The rate of success for detecting a ROI for each subject was as follows: lower left visual quadrant 96 percent (48 out of 50 subjects), lower right visual quadrant 100 percent (50 out of 50 subjects), upper left visual quadrant 100 percent (50 out of 50 subjects), upper right visual quadrant 92 percent (46 out of 50 subjects). As with the PMs at the group level, peak vertex distribution maps at the single-subject level (Figure 7) showed less spatial variability for CBA compared to SBAV. Furthermore, for CBA compared to SBAV we observed an increase in the number of multiple overlapping single-subject ROI peak vertices per vertex for each visual quadrant and a corresponding decrease in the number of single overlapping single-subject ROI peak vertices per vertex (Table 6).

**Table 6.**
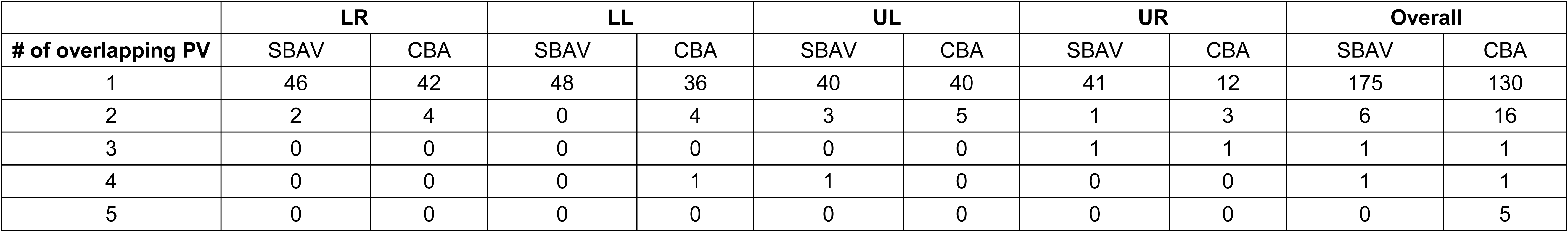
Single-subject ROI peak vertex distribution mapping. Number of overlapping single-subject peak vertices for each visual quadrant before and after macroanatomical alignment (SBAV and CBA). Overall, there was an increase in the number of multiple overlapping single-subject ROI peak vertices per vertex after macroanatomical alignment (CBA) for each visual quadrant and a corresponding decrease in the number of single overlapping single-subject ROI peak vertices per vertex. PV = ROI peak vertex.

## DISCUSSION

The aim of our study was to evaluate the utility of CBA for an attention-enhanced visual field localizer paradigm used to map a circumscribed region within retinotopically organized visual areas. Our paradigm mapped homologous regions in each visual quadrant reliably across early visual areas. As expected, CBA led to a marked reduction in macroanatomical variability with a number of beneficial effects on the functional level, which clearly exceeded those observed for SBAV.

Compared to VBA and SBAV, CBA resulted in the largest improvement in the group ROI analysis across visual quadrants (Figure 4). This was reflected in the consistent size increase of the resulting group ROIs compared to VBA and SBAV (Table 1), which indicates an increased power to detect subregions of early visual areas show-ing position selectivity. Unsurprisingly, we did not observe an increase of position selectivity when comparing visual quadrant ROIs across alignment techniques (Table 2). Importantly, only CBA but not SBAV preserved the vertical symmetry of group ROIs characteristic for early visual areas, which was already evident for VBA (Table 3).

Regarding changes in the probability of activation overlap across the three alignment techniques as reflected in the PMs, a clear pattern emerged. The probability of activation overlap increased gradually with each step, peaking for CBA with a value of about 75 percent. For SBAV, effects were weaker and considerably more wide-spread, favoring more peripheral brain regions. Similarly, PDMs showed an increase in the probability of activation overlap of up to 40 percent in the central region of interest for CBA, resulting in considerably more focused activation patterns, while the opposite effect emerged in more peripheral vertices (Figure 6). This is most likely not attributable to a decreased spatial overlap in the periphery of early visual areas. Rather, it demonstrates that CBA consistently reduces spurious spread-out activation resulting from poor macroanatomical correspondence after VBA and a generalized increase in the SNR due to SBAV. It also suggests that VBA and SBAV might misrepresent the location and extent of early visual areas. This is notion is supported by changes of the center gravity of group ROIs between SBAV and CBA, which was particularly pronounced in the lower left visual quadrant (Table 1). Together, these findings indicate that CBA substantially increases statistical power when studying early visual areas at the group level. Naturally, this effect of CBA should also extend to studies with a more global focus, such as connectivity analyses^35, 55^.

The specific advantages of CBA compared to SBAV were also evident in the markedly decreased variability of single-subject ROI peak vertex locations for each visual quadrant (Figure 7, Table 6). This indicates that the reduction of macroanatomical and functional inter-subject variability achieved by CBA is the main reason for our improved group-level results. Our findings confirm the notion that transforming functional data from volume-space into surface space already increases statistical power by reducing signal contamination from non-neuronal tissue, thus improving the signal-to-noise ratio. Therefore, using SBAV as a proxy for VBA would underestimate the actual benefits of CBA, which clearly exceed those of SBAV. Overall, the combined beneficial effects of an improved signal-to-noise ratio and reduced macroanatomical alignment inherent in the CBA approach and evident in our data support the notion that CBA is clearly the superior alignment technique when studying the visual system.

For VBA we observed the largest group ROIs for the right and left lower visual quadrant, an effect that changed after SBAV and CBA (Figure 4a, Table 1). For SBAV we observed the largest group ROIs for the left upper and right lower visual quadrant, which did not persist after CBA (Figure 4b & 4c, Table 1). Notably, several studies using volume-based fMRI and magnetoencephalographic analyses reported lateralized effects on neurophysiological parameters in early visual areas^56, 57^. Our observation raises the question, whether these findings could at least partly be explained by lateralized differences in macro-anatomical variability rather than true functional differences.

Conversely, our CBA-aided group analysis allowed us to compare the response properties of each visual quadrant in a more unbiased way. We observed larger group ROIs for the lower visual hemifield. In a CBA-based probabilistic atlas of the visual system, which included all regions that could be defined in more than fifty percent of subjects, probabilistic ROIs for dorsal V1 and V2 were also noticeably larger probabilistic ROIs for ventral V1 and V2, whereas this effect was less clear for V3^44^. These results are in line with our own findings and could be attributable to higher residual anatomical variability after CBA in ventral occipital cortex representing the upper visual hemifield. Alternatively, they could be due to true differences in response properties such as receptive field size or overall area size. The latter interpretation is supported by studies showing functional differences between upper and lower visual hemifields already at the retinal level in the form of differences in receptor densities. Cone density was higher in the superior parts of the retina, which processes information from lower visual fields. Conversely, higher rod density was observed in the inferior parts^58, 59^. Moreover, Eickhoff et al. observed dorso-ventral asymmetries in receptor densities in V2 and V3^58^ and reported a higher density of GABA-A, and muscarinic M3-receptors in ventral parts of V2 and V3. Furthermore, there is evidence for fundamental differences in receptive field shape from a pRF mapping study. Estimating both the aspect ratios and the size of the mapped areas, a more elliptical receptive field shape was observed for the upper visual hemifield represented by ventral parts of the visual cortex being compared to the lower visual hemifield represented by dorsal parts of the visual cortex^60^. Additionally, there is evidence for a behavioral advantage in the lower visual hemifield for shape discrimination as well as higher BOLD-signal changes and peak amplitudes of MEG/EEG responses^51, 53, 54, 61, 62^. Together, these findings demonstrate clear differences in the functional architecture of early visual areas representing the upper and lower visual hemifield. This has been attributed to the fact that the lower visual hemifield represented by dorsal parts of the occipital lobe is more closely linked to the dorsal visual pathway, while the upper visual hemifield represented by ventral parts of the occipital lobe is more closely linked to the ventral visual pathway^63, 64^. Furthermore, there is evidence for fundamental differences in receptive field shape from a pRF mapping study. For the upper visual hemifield represented by ventral parts of the visual cortex an increased size and more elliptical shape of receptive fields was observed compared to the lower visual hemifield represented by dorsal parts of the visual cortex^60^. This implies, that the lower visual field is more specialized for the precise localization and representation of space. Our observation of larger ROIs in the lower visual hemifield is in line with these findings. Thus, our results imply that CBA is a useful tool to extend the study of functional and behavioral asymmetries in early visual areas to the group level.

One important limitation of the current study is the lack of a complementary retinotopic mapping data set due to time constraints. This data would have allowed us to delineate the boundaries of early visual areas and pinpoint the exact visual area containing each individual single-subject ROI. Retinotopic mapping studies indicate that peak activation of single subjects assessed by visual localizers are not consistently located in the same visual area. Most localizer tasks show peak activation not in V1 but rather in V2 or V3^12^. It is therefore highly likely that our single-subject peak activation did not consistently belong to the same visual cortical area. With the current data set we cannot determine how precisely individual visual areas were aligned with CBA, and whether individual levels of the visual cortical hierarchy were differentially affected. However, the position of our group ROIs, which bordered the calcarine sulcus and spanned the occipital pole, indicate that they mainly comprised V2 and V3.

Similarly, after CBA a comparable increase in the probability of overlap was observed in the same part of occipital cortex. While this is at least suggestive of a relatively consistent benefit of CBA across visual areas, more fine-grained studies including retinotopic mapping are required to definitively address this question.

Furthermore, we did not use eye tracking to insure sufficient fixation. We also did not include an additional central attentional control task on the fixation cross, which would have further encouraged continuous fixation. This was done deliberately in order to keep the difficulty level adequate for psychiatric patient populations. Our average success rate for finding reliable activation in early visual areas across all four visual quadrants was 97 (92 - 100) percent. Insufficient fixation might explain our failure to find reliable activation in a small fraction of subjects in the upper right and lower left visual quadrant.

Finally, several properties of the VBA data set differed from the SBAV and CBA data sets. We could not match volume-based and surface-based preprocessing parameters completely due to inherent differences between the three-dimensional and two-dimensional spatial smoothing algorithms employed. The number of voxels and vertices containing functional data was not identical, differentially affecting Bonferroni correction of group results. However, the smaller analysis space of the VBA data set, 69 percent the size of the SBAV and CBA data set, should only underestimate the effect of the additional processing steps featured in the SBAV and CBA analysis.

Our study also has implications beyond mapping the visual system in healthy populations. Visual processing deficits are a prominent feature of neurodevelopmental psychiatric disorders such as ADHD, schizophrenia and autism spectrum disorders^7, 8, 65– 72^, which can also perturb crucial higher order cognitive processes including working memory^73–75^. The current localizer paradigm will be useful to investigate local impairments of visual information processing as well as disturbances in the interplay between early visual areas and brain networks supporting higher-order cognitive processes. Here, CBA will be particularly relevant to reduce the confounding effects of increased macroanatomical variability in disorders such as schizophrenia in order to measure true group differences and true functional variability^38, 76^. On the other hand, CBA might also be crucial for investigating the neurodevelopmental underpinnings of increased macroanatomical variability itself. To this end, the inclusion of probabilistic atlases containing information about gene expression profiles^77^ as well as cyto and receptor architectonics^78, 79^ will be valuable.

Our CBA approach relied solely on cortical curvature information to reduce macroanatomical variability. The advantage of this method is its feasibility for the vast majority of fMRI data sets, since it only requires a structural brain scan of sufficient quality and resolution. Among comparable methods, it is the most data driven and objective approach. However, the achievable reduction of macroanatomical variability is limited by the variable and imperfect correlation between brain structure and brain function^34, 40^. Therefore, more advanced methods additionally utilize orthogonal functional data to further reduce anatomical variability, including the use of functional activation or connectivity patterns to improve macroanatomical alignment across the whole brain^20, 80, 81^. Additionally, a more complex approach has been proposed, which aligns cortical data using ‘areal features’ more closely tied to cortical areas than cortical folding patterns, including maps of relative myelin content and functional resting state networks^82^. These methods have shown to provide a relevant additional reduction of macroanatomical variability for a variety of paradigms including visual functional localizers. Future studies should also evaluate these methods for retinotopic mapping and visual field localizers. Moreover, it has been demonstrated for early auditory areas, that the additional use of a probabilistic atlas of cytoarchitectonically defined areas further enhance standard CBA results^83^. In principle, such an approach would easily be feasible for the visual system.

To summarize, we demonstrated the clear advantages of CBA compared to VBA for the analysis of visual field localizer data at the group level, signified by a considerable reduction of spatial variability across subjects of up to forty percent across early visual areas. Our findings extend previous CBA studies examining other major categories of visual mapping techniques. They underscore the loss of information and statistical power incurred by the use of VBA methods in the majority of fMRI studies.

Therefore, CBA and comparable methods should be seriously considered as a standard procedure for the detailed study of visual information processing and its disturbance in mental disorders.

## METHODS AND MATERIALS

### Participants

All participants gave their written informed consent to participate in the study in accordance with the study protocol approved by the ethical review board of the Faculty of Medicine at Goethe University. The experimental procedures were conducted in conformity with the approved guidelines and the Declaration of Helsinki. Individuals received compensation for their participation. We recruited 51 healthy volunteers (female : male = 28 : 23) with age ranging between 18-43 years (mean = 24). All participants were non-smokers, had no history of neurological or psychiatric illness and reported normal or corrected-to-normal visual acuity. One participant was left-handed as assessed by the German version of the Edinburgh Handedness Inventory^84^.

### Stimuli and task

Subjects performed a visual field localizer paradigm (Figure 1a) implemented using Presentation (Neurobehavioral Systems, Version 18.0) as part of a larger study investigating the role of visual areas for higher cognitive functions. The task consisted of a series of flickering black-and-white-colored round shaped checkerboard stimuli (flicker frequency = 7.5 Hz). Checkerboard stimuli appeared for 2000 ms randomly at one of four different locations (non-target trial). Each location mapped a homologous position in one of the four visual quadrants. The regular inter-trial interval (ITI) was 0 ms. However, once every 10 to 14 trials (11 times overall), the ITI increased to 2000 ms (prolonged ITI). Our paradigm featured a simple target-detection task. During 36 trials, the two centrally located squares of the checkerboard changed their color to yellow for 133 ms (target trial). Participants were instructed to press a response box button with their left thumb as quickly as possible if they detected a target. The task consisted of a total of 144 trials (36 target trials, 108 standard trials). This target probability of 25 percent resulted in one target trial every fourth trial on average (range 3 to 5 trials) (Figure 1a). Throughout the task a black, x-shaped fixation cross was displayed at the center of the screen. Participants were instructed to continuously fixate on the fixation cross. Before the first trial, only the fixation cross was displayed for 10 seconds. After the last trial, only the fixation cross was displayed for 20 seconds. The total duration of the paradigm was 340 seconds (Figure 1b). For the purpose of our analyses we defined a total of four conditions, one for each of the four stimulus locations. Each participant practiced the task prior to the measurement.

**Figure 1.**
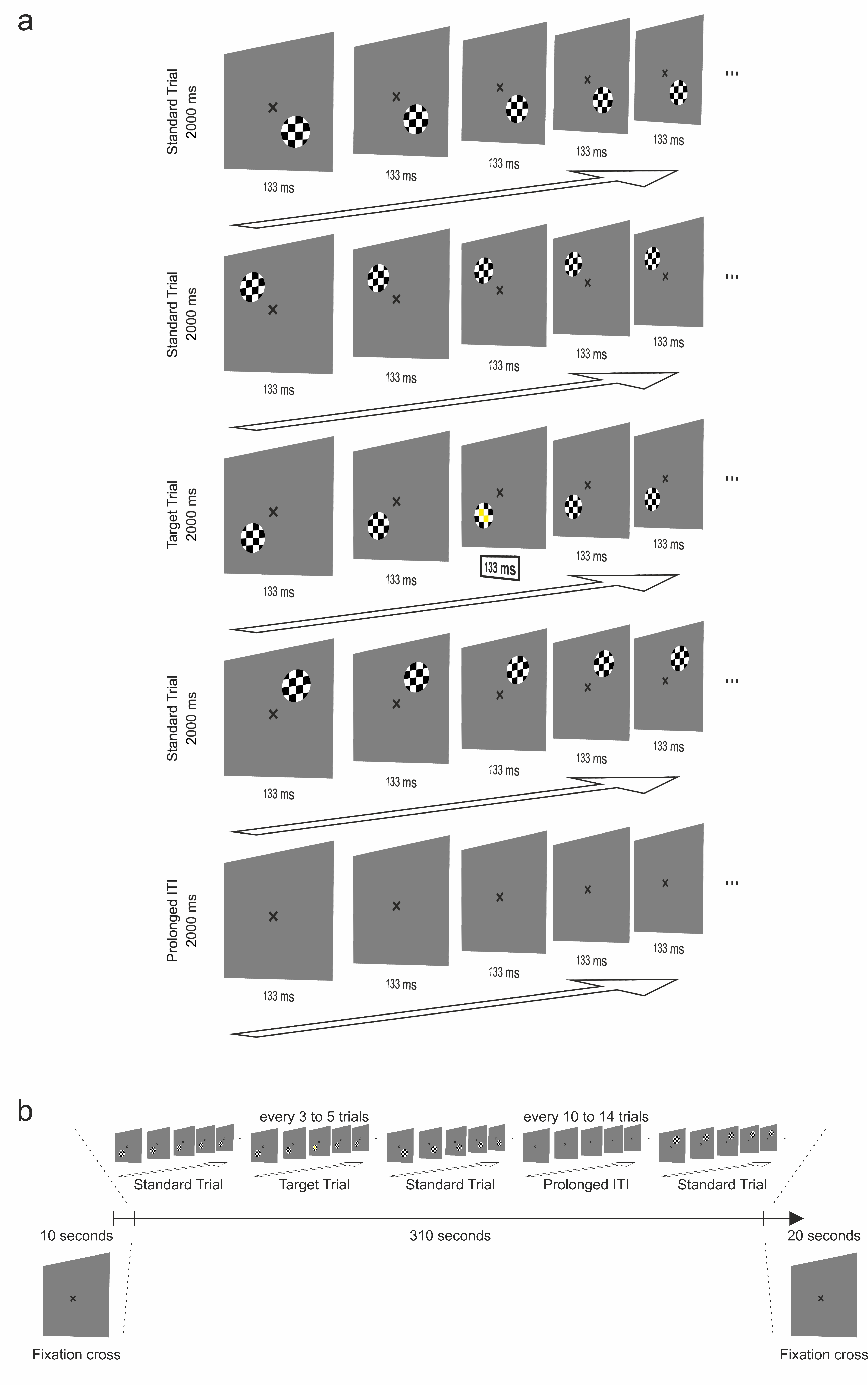
Visual field Localizer Paradigm. (a) The task consisted of flickering, black-and-white colored checkerboards that appeared randomly at homologous positions of the participant’s visual quadrant. Only in 25 percent of the trials the two centrally located squares changed their color into yellow for 133 ms. Participants were required to press a response box button when noticing that. During the whole task participants were instructed to fixate a black, x-shaped fixation cross presented in the center of the screen. Checkerboards appeared for 2000 ms. The regular inter-stimulus interval was 0 ms. (b) Every 10 to 14 trials, the ITI extended to 2000 ms. The task comprised 144 trials (25 percent target trials). It was preceded and followed by a presentation of the fixation cross for 10 seconds.

### Acquisition and analysis of fMRI data

We acquired functional MRI data on a Siemens 3T MAGNETOM Trio scanner at the Goethe University Brain Imaging Centre using a gradient-echo 2D EPI sequence (32 axial slices, TR = 2000 ms, TE = 30 ms, FA = 90°, FoV = 192 x 192 mm2, voxel size = 3 x 3 x 3 mm^3^, gap = 1 mm, effective slice thickness = 4 mm). Slices were positioned parallel to the anterior- and posterior commissure. Functional images were acquired in a single run comprising the acquisition of 170 volumes. Immediately before each functional run, 6 volumes of this 2D EPI sequence were acquired with identical parameters except for a switch of phase encoding direction (posterior to anterior instead of anterior to posterior) for EPI distortion correction. Anatomical MRI data for cortex reconstruction and co-registration with functional MRI data was acquired with a high-resolution T1-weighted 3D volume using a Magnetization-Prepared Rapid Gradient-Echo (MP-RAGE) sequence (192 sagittal slices, TR = 1900 ms, TE = 3.04 ms, TI 900 ms, FA = 9°, FoV = 256 x 256 mm^2^, voxel size = 1 x 1 x 1 mm^3^). Stimulus presentation was constantly synchronized with the fMRI sequence. Head motion was minimized with pillows. The task was projected by a beamer onto a mirror attached on the head coil. MRI data were pre-processed and analyzed using BrainVoyager 20.6^85^, the NeuroElf Matlab toolbox (www.neuroelf.net) and custom software written in Matlab. One subject had to be excluded due to excessive intra-scan motion.

### Structural image pre-processing

Structural data pre-processing included background cleaning, brain extraction and bias field correction to minimize image intensity inhomogeneities^85^. Bias field correction employed a “surface fitting” approach using singular value decomposition based least squares low-order (Legendre) polynomials to model low-frequency variations across 3D image space^86^. We used polynomials with an order of three, which were fitted to a subset of voxels labeled as belonging to white matter. The estimated parameters of the polynomials were used to construct a bias field which was removed from the data. Our approach comprised of one iteration using automatic white matter labeling^87^ and four iterations using manual white matter labeling.

Subsequently, structural data were transformed into Talairach coordinate space^29^. This comprised manual labeling of the anterior commissure (AC) and posterior commissure (PC) as well as the borders of the cerebrum. These landmarks were then used to rotate each brain in the AC-PC plane followed by piece-wise, linear transformations to fit each brain in the common Talairach “proportional grid” system^19^. Transformation into Talairach coordinate space was performed because the subsequent automatic segmentation procedure exploits anatomical knowledge for initial brain segmentation including removing subcortical structures and disconnection of cortical hemispheres. To prepare the data for this procedure, we performed a manual filling of the lateral ventricles. Based on the automatic segmentation of the structural scans along the white-gray matter boundary^88^, cortical hemispheres were reconstructed into folded, topologically correct folded mesh representations, which were tessellated to produce surface reconstructions and calculate curvature maps reflecting the individual cortical folding patterns. Surface reconstructions were subsequently morphed into distortion corrected spherical representations. Finally, both folded and spherical mesh representations were downsampled to a standard number of vertices (40962 vertices per hemisphere, mean vertex distance: 1.5 mm). We used these standardized mesh representations for all surface-based processing steps.

### Cortex-based alignment of structural data

We then applied for each hemisphere separately a high-resolution, multiscale cortex-based alignment procedure based on the individual curvature maps of all 50 participants. This CBA approach, which reliably aligns corresponding gyri and sulci across subjects^85^, consists of an initial rigid and a subsequent non-rigid alignment step^19^ (Figure 2a & 2b). During the initial step, cortical folding patterns of each sphere are aligned rigidly to the cortical folding pattern of a single target sphere by global rotation. Rigid CBA operates only on highly smoothed curvature maps containing only the most prominent anatomical landmarks. We used the rotation parameters with the highest degree of overlap between the curvature of each individual sphere and the target sphere as the starting point for the subsequent non-rigid CBA.

**Figure 2.**

CBA consisted of a rigid alignment to a single target brain and a non-linear alignment to an iteratively updated group average brain. **(a)** We carried out an initial CBA solely to generate an unbiased average target brain for the final CBA. A randomly selected brain from among all participants was used for the initial rigid CBA. **(b)** For the final CBA we used the unbiased average target brain created during the initial CBA for rigid CBA. **(c)** We generated average surface representations before and after macroanatomical alignment for each hemisphere, which we subsequently merged, inflated and used for analysis and visualization of the appropriate data sets.

Non-rigid CBA employs a coarse-to-fine matching strategy, which operates sequentially at four levels of curvature smoothing starting with the detail level used during rigid CBA. Each subsequent level includes increasingly finer anatomical details up to almost the full curvature information. Importantly, non-rigid CBA aligns each cortical folding pattern to a dynamically updated group average through iterative morphing. This moving target approach, which generates the target curvature map from the average curvature across all hemispheres at a given alignment stage avoids the possible confounding effects of a suboptimal selection of an individual target brain, whose folding pattern might deviate considerably from the cohort average.

Notably, rigid CBA typically utilizes a single brain randomly drawn from the cohort to be aligned as its target brain. However, the folding pattern of this brain might also deviate considerably from the cohort average. To also address this potential confound, we first conducted a preliminary CBA encompassing both rigid and non-rigid macro anatomical alignment (Figure 2a). We then conducted a second, final CBA. Here, we used the aligned average brain derived from the preliminary CBA as an unbiased target for the rigid alignment step (Figure 2b). After the final non-rigid CBA, we merged both hemispheres of each individual brain to create a global surface-based analysis space.

Furthermore, for each hemisphere we created average surface representations from the original, non-aligned folded mesh representations, which were subsequently merged, inflated and used for data analysis and visualization. We repeated these steps after applying the transformation matrix of the final rigid and non-rigid CBA to the folded mesh representations, yielding an accurate representation of the structural effects of macroanatomical alignment (Figure 2c).

### Functional image pre-processing

The first four volumes of each functional run were discarded to allow for T1 equilibration. Initial volume-based pre-processing of functional MRI data comprised slice timing correction using sinc interpolation and 3D motion correction using sinc interpolation. Next, we performed echo-planar imaging distortion correction using the Correction based on Opposite Phase Encoding method^89, 90^. EPI distortion corrected functional data were co-registered to the untransformed extracted brains. This was accomplished utilizing a boundary-based registration algorithm optimized for surface-based analyses^91^. After co-registration to the fully cleaned but untransformed structural data, functional data were transformed into Talairach coordinate space by applying the transformation matrix generated during Talairach transformation of the anatomical data using sinc interpolation. This transformation preserved the original voxel size of the functional data (3 x 3 x 3 mm^3^) (Figure 3).

**Figure 3.**
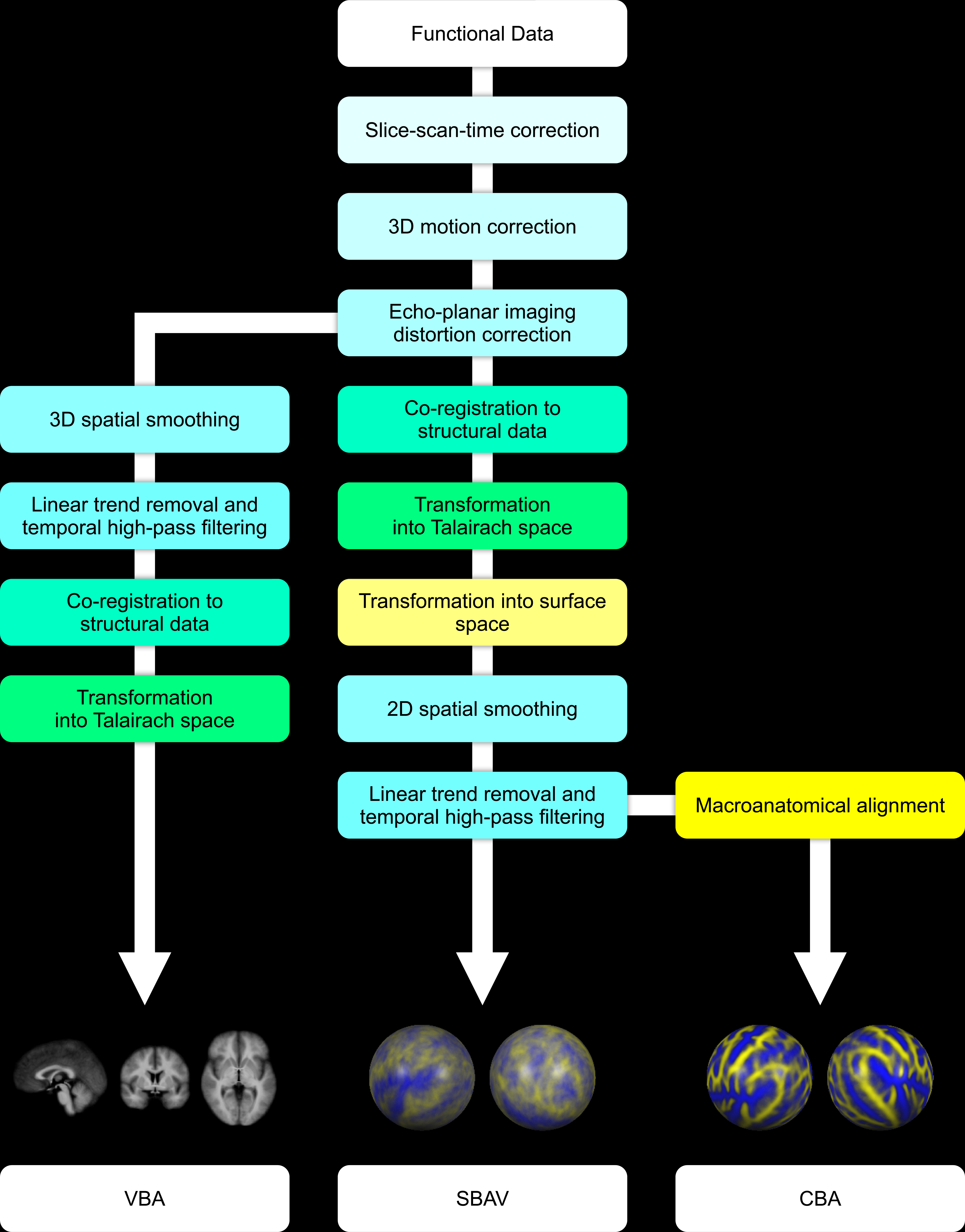
Sequences of functional data pre-processing, coregistration of structural and functional data and spatial transformation operations used to generate the three functional data sets used in our study: VBA, SBAV and CBA. For VBA we conducted all data pre-processing operations in volume space, including slice-scan-time correction, 3D motion correction, echo-planar imaging distortion correction, 3D spatial smoothing and linear trend removal with temporal high-pass filtering. Finally, functional data were co-registered to the structural data and transformed into Talairach space. For SBAV and CBA, we conducted all data pre-processing operations up to echo-planar imaging distortion correction in volume space. Here, co-registration of functional data to the structural data and transformation into Talairach space was followed by transformation into surface space. We then conducted 2D spatial smoothing and linear trend removal with temporal high-pass filtering in surface space. For CBA only, we then applied macroanatomical alignment.

**Figure 4.**
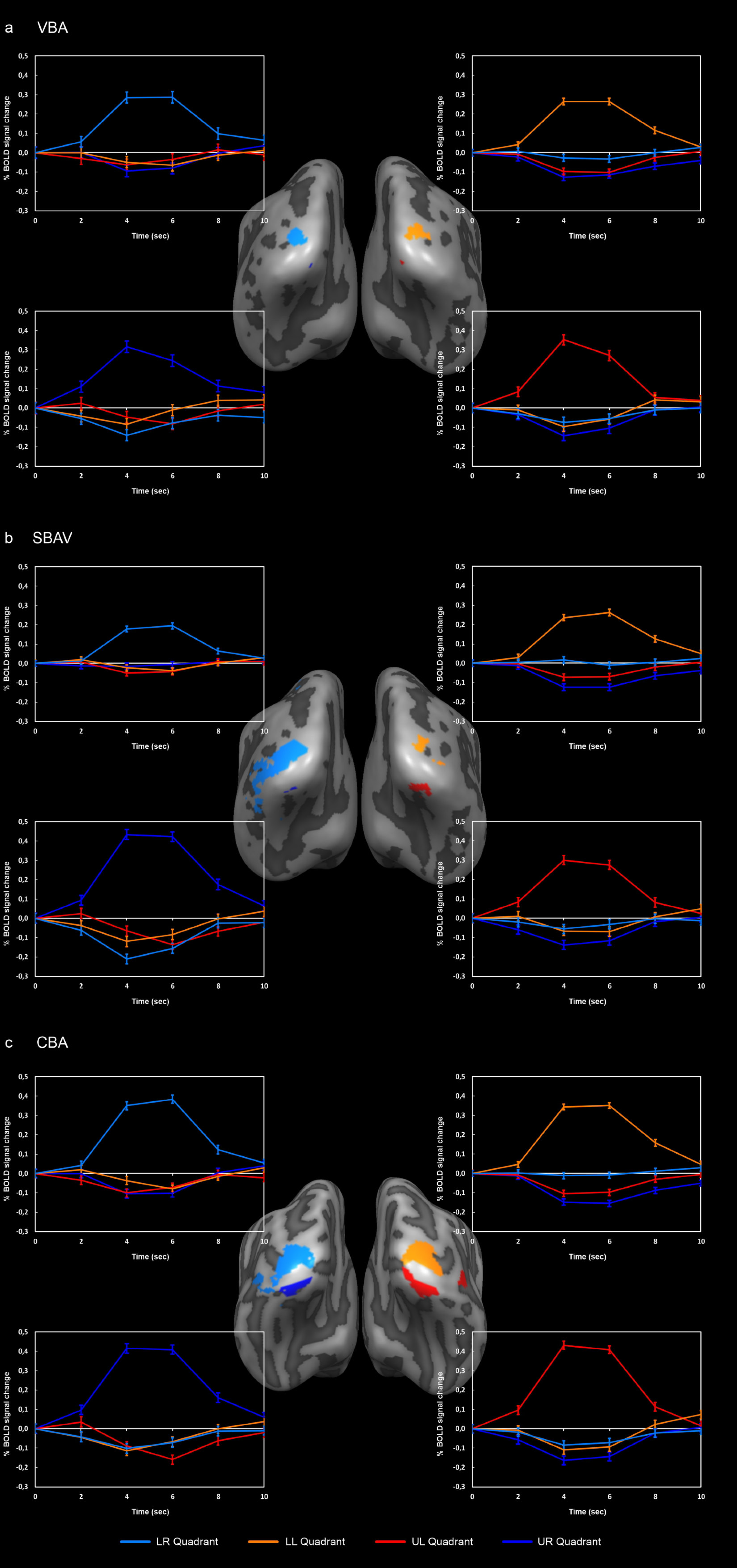
Group analysis of visual quadrants. **(a)** VBA results. Maps and average timecourses were computed in volume space; maps were projected on the non-aligned average surface representation. **(b)** SBAV results. Maps and average timecourses were computed in surface space; maps were projected on the non-aligned surface representation. **(c)** CBA results. Maps and average timecourses were computed in surface space; maps were projected on the aligned average surface representation. Overall, three out of four visual quadrant ROIs exhibited a pattern of continuously increasing cluster size reflecting an increasing extent of significant position selectivity across alignment techniques. Conversely, the ROI of the lower right visual quadrant showed a cluster size decrease after CBA. Average timecourses (incl. SEM) showed clear position selectivity with a strong BOLD signal increase for the position of interest and no BOLD signal increase for the other three positions. ROI/graph colors: light-blue = lower right (LR) visual quadrant, orange = lower left (LL) visual quadrant, red = upper left (UL) visual quadrant, dark-blue = upper right (UR) visual quadrant.

**Figure 5.**

Probability maps (PMs). PMs indicating the probability of activation overlap across subjects for each visual quadrant. The color code gray-to-white indicates the probability of activation overlap of single-subject maps thresholded at a minimum of ten percent probability of activation overlap. Single-subject maps were thresholded at p<0.05 (uncorr.). PMs were thresholded at a minimum of ten percent probability of activation overlap. We also applied a cluster level threshold of 100 vertices. **(a)** PMs for VBA showed a maximum probability of activation overlap of up to 50 percent. **(b)** PMs for SBAV showed a maximum probability of activation overlap of up to 45 percent. **(c)** PMs for CBA showed a maximum probability of activation overlap of up to 75 percent.

**Figure 6.**

Probability Difference Maps (PDMs). PDMs indicating the differential impact of the individual steps of our overall macroanatomical alignment approach for each visual quadrant. PDMs were generated using PMs derived from single-subject maps thresholded at minimum of zero percent probability. The color code indicates the difference of activation overlap. The color code brown-to-white indicates a higher degree of functional activation overlap for the more advanced alignment method. The color code blue-to-green indicates a higher degree of functional activation overlap for the less advanced alignment method. PDMs were thresholded at a minimum probability difference of five percent. **(a)** The impact of surface-based functional data readout and pre-processing compared to standard volume-based alignment (SBAV minus VBA) was characterized by a widespread activation with an increase in the probability of activation overlap of up to 15 percent around the central ROIs and a decrease in the probability of activation overlap of up to 10 at the location of the central ROIs. **(b)** The additional impact of macroanatomical alignment (CBA minus SBAV) was less widespread but characterized by an increase in the probability of activation overlap of up to 35 percent at the location of the central ROIs and a decrease in the probability of activation overlap of up to 35 percent around the central ROIs. **(c)** The additive impact of both methodological elements (CBA minus VBA) was characterized by an increase in the probability of activation overlap of up to 30 percent at the location of the central ROIs and a decrease in the probability of activation overlap of up to 20 percent around the central ROIs.

**Figure 7.**
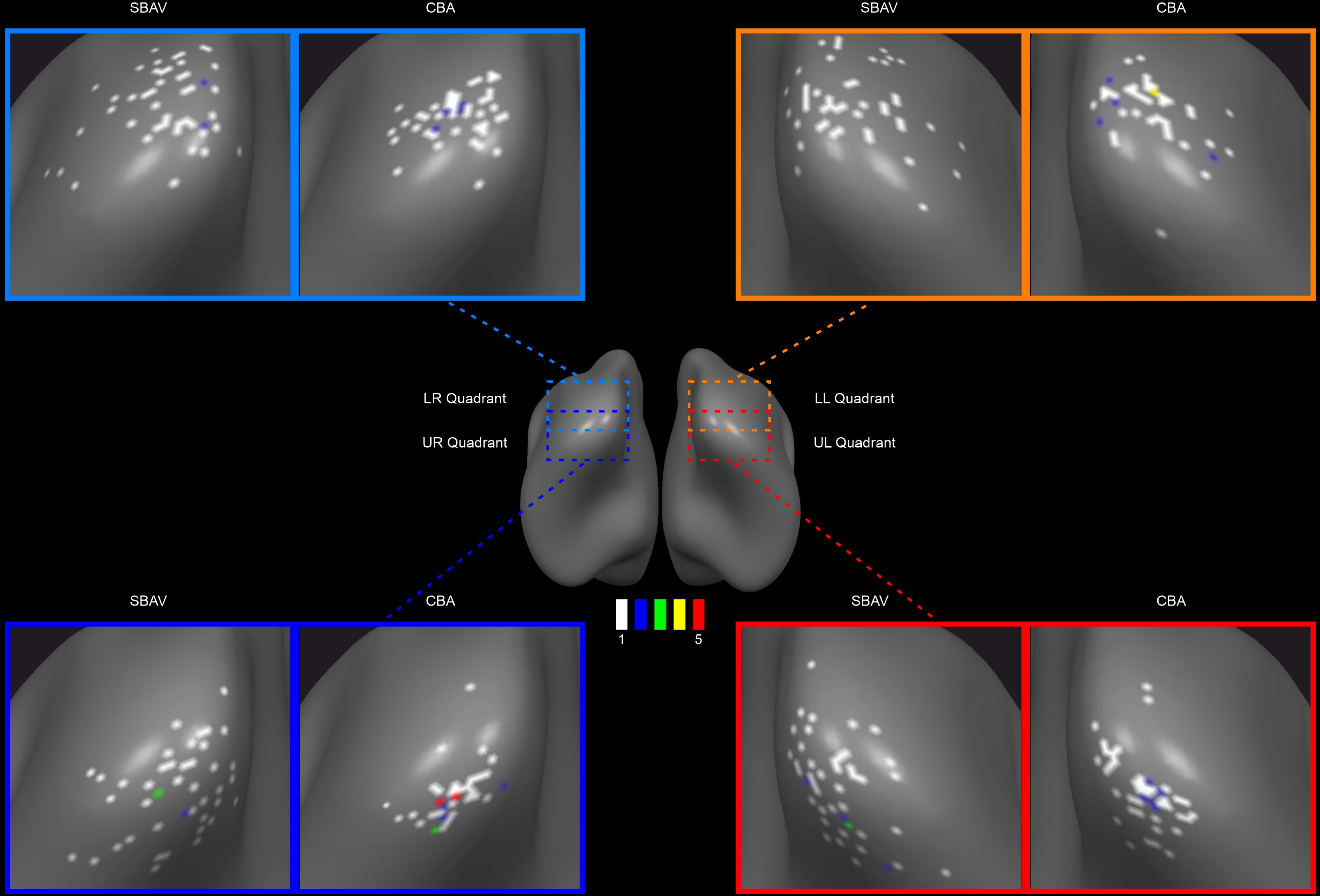
Single-subject peak vertex distribution maps for SBAV and CBA data sets. We mapped single-subject peak vertices for each visual quadrant in surface space for SBAV and CBA data. We then computed the vertex-wise number of single-subject peak vertices. The color code indicates the number of overlapping single-subject peak vertices per vertex. An increase in the number of overlapping single-subject ROI peak vertices per vertex was observed after macroanatomical alignment (CBA). For each visual quadrant, single subject peak vertices were mapped in surface space and the vertex-wise number of single subject peak vertices was computed before and after macroanatomical alignment. The color code indicates the number of overlapping single-subject peak vertices per vertex. A nominal increase in the degree of overlap single subject peak vertices was observed after macroanatomical alignment (CBA).

### Surface-based pre-processing

The volumetric functional data were then transformed into surface space by sampling on the individual cortical surface reconstructions incorporating data from −1 to +3 mm along vertex normals using trilinear interpolation. Subsequent pre-processing of functional MRI data in surface space started with spatial smoothing using a nearest neighbor interpolation (1 iteration). Based on the standardized vertex distance of 1.5 mm this approximates a 2D Gaussian smoothing kernel with a full width at half maximum (FWHM) of 3 mm. We opted for minimal spatial smoothing to prevent a loss of accuracy for our visual field localizer. Spatial smoothing was followed by linear trend removal and temporal high-pass filtering using fast Fourier transformation (high-pass 0.00903 Hz). Based on the vertex-to-vertex referencing from the folded, topologically correct surface reconstructions to the spherical representations, we mapped the fully preprocessed functional data into a common spherical coordinate system (Figure 3).

Finally, we applied surface-based anatomical masks that only included cortical vertices in our analysis to the functional data. These masks excluded those subcortical structures, i.e., parts of thalamus and the basal ganglia, which mapped onto the midline of our surface reconstructions. For functional data analysis and subsequent Bonferroni correction in surface space, this yielded a total number of 76132 vertices.

### Full volume-based pre-processing

To generate a purely volumetric data set for the comparison of VBA and SBAV, preprocessing after EPI distortion correction was also conducted in volume space mirroring as closely as possible the steps and parameters outlined above for surface-based pre-processing. First, we applied spatial smoothing using a 3D Gaussian smoothing kernel with a FWHM of 3 mm, which approximates the degree of surface-based spatial smoothing. Second, we applied linear trend removal and temporal high-pass filtering using fast Fourier transformation (high-pass 0.00903 Hz) using parameters exactly matching surface-based pre-processing (Figure 3). This data set was not transformed into surface space and did not include an anatomical mask. For functional data analysis and subsequent Bonferroni correction in volume space, this yielded a total number of 52504 voxels. Thus, the analysis space for VBA was 69 percent the size of the analysis space for SBAV and CBA (52504 voxels vs. 76132 vertices).

### Final functional data sets

Overall, we generated three different functional data sets: a volume-based data set, which was fully pre-processed and aligned in volume-space (VBA); a surface-based data set, for which spatial smoothing and temporal filtering were only applied after transformation in surface space, but without macroanatomical alignment (SBAV); a surface-based data set, which was pre-processed in the same way as the SBAV data set and also utilized macroanatomical alignment (CBA) (Figure 3). Accordingly, primary analysis of these datasets was carried out in volume space (VBA) and surface space (SBAV, CBA) respectively.

Planned direct comparisons between these three data sets allowed us to evaluate the effects of different steps of our macroanatomical alignment approach. We compared the VBA and SBAV data sets to assess in isolation the impact of surface-based preprocessing while keeping anatomical alignment constant. We compared the SBAV and CBA data sets to assess in isolation the impact of macroanatomical alignment while keeping preprocessing parameters constant. Finally, we compared the VBA and CBA data sets to assess the combined impact of both surface-based preprocessing and macroanatomical alignment.

### fMRI group analysis of visual quadrants

We performed multi-subject statistical analyses using multiple linear regression of the BOLD signal. The presentation of each checkerboard stimulus sequence at a single location, was modelled by an ideal box-car function, which covered the volume of each trial, convolved with a synthetic two-gamma function. These predictors were used to build the design matrix of the experiment. Individual statistical maps were generated by associating each voxel with the beta-value corresponding to the specific set of predictors and calculated on the basis of the least mean squares solution of the general linear model. The resulting individual statistical maps were entered into a second-level random-effects group analysis using a summary statistic approach.

We performed analyses focusing on the mapping of the four visual quadrants at the group level. To define the group-level ROI for each visual quadrant, we computed separate weighted contrasts for each quadrant against the other three quadrants. We assigned a weight of three to the position of interest, e.g. (β*^Quad^*^1^ x 3) / (β*^Quad^*^2^ + β*^Quad^*^3^ + β*^Quad4^*) (p < 0.05, Bonferroni corrected). This allowed us to detect brain regions showing significant position selectivity. For each resulting group-level ROI, we extracted average time courses (incl. standard errors of the mean = SEM) for all four conditions. We conducted this analysis for all three data sets (VBA, SBAV, CBA). For the VBA data set, we computed this analysis completely in volume space using the original resolution of the functional data (voxel size: 3 x 3 x 3 mm^3^). We projected the resulting maps on the non-aligned average surface representation to achieve an appropriate visualization and to allow a comparable quantification of cluster size for the resulting ROIs, i.e., vertices instead of voxels. To this end, volumetric functional maps were transformed into surface space by sampling on the average cortical surface incorporating data from −1 to +3 mm along vertex normals of the group average surface brain using trilinear interpolation.

We used two approaches to determine, whether position selectivity differed between our three data sets: First, to assess differences in the extent of position selectivity across early visual areas, we compared the ROI size, i.e. the number of vertices, for each position of interest across data sets. Second, to test for differences in the strength of position selectivity within corresponding ROIs we conducted separate linear mixed models with random intercept for each visual quadrant using the t-values of each subject as the dependent variable and the alignment methods (VBA, SBAV and CBA) as the independent variable. To correct for multiple comparisons, p-values were adjusted using Bonferroni correction.

Finally, to assess the impact of the three alignment approaches on horizontal and vertical symmetry of our group-level ROIs, we computed an asymmetry index (AI) based on ROI size, i.e the number of vertices, following an established procedure^92^, between each pair of ROIs using the following formula: (| size*^ROI1^* - size*^ROI2^* | / size*^ROI1^* + size*^ROI2^*) * 100.

### Probability maps

To quantify and visualize variability of functional activation and possible changes due to macroanatomical alignment, the use of PMs has been proposed. PMs are specifically useful to assess inconsistencies, i.e., disparities between individuals regarding the location of a particular (visual) area^93, 94^. To quantify the spatial consistency of position selective activation patterns, we generated PMs for each visual quadrant for all three data sets (VBA, SBAV, CBA). These maps represent the relative number of subjects showing significant task activity in our single-subject analysis. To this end, we generated single-subject t-maps based on the same weighted contrasts employed in the group analysis but set at a more lenient statistical threshold (p < 0.05 uncorrected). PMs were calculated by counting the number of subjects showing above-threshold activation in their individual t-maps at a given vertex, dividing this value by the total number of subjects, and multiplying the result by 100. For the VBA data sets, we computed all of these steps in volume space and transformed the final PM into surface space using the same parameters outlined above for the volumetric group maps. Finally, all PMs were thresholded at a minimum of ten percent probability of activation overlap. We also applied a cluster level threshold of 100 vertices to focus on the main areas of interest, i.e., the visual quadrants. Additionally, we counted the number of vertices in the corresponding probability maps exceeding the threshold of ten percent probability of activation overlap for each visual quadrant and analysis methods. Our goal was to quantify and compare the extent of early visual cortex, where each analysis method had a relevant impact on the probability of activation overlap.

### Probability difference maps

Additionally, we aimed to quantify changes in spatial consistency of position selective activation patterns as a result of the different alignment methods. To this end, using the original unthresholded PMs we calculated PDMs for each visual quadrant, thresholded at a minimum probability difference of five percent. The resulting three PDMs capture different aspects of our overall approach: the impact of surface-based functional data readout and pre-processing compared to volume-based alignment (SBAV minus VBA), the additional impact of applying macroanatomical alignment (CBA minus SBAV) and the additive impact of both methods (CBA minus VBA). Additionally, we counted the number of vertices in the corresponding PDMs exceeding the threshold of 5 percent difference in probability of activation overlap for each visual quadrant. Our goal was to quantify and compare the extent of early visual cortex, where we observed a difference in the probability of activation overlap for a comparison of analysis method.

### Single-subject ROI peak vertex distribution mapping

For single-subject level analyses, we first defined ROIs for each subject independently before and after macroanatomical alignment, i.e., for SBAV and CBA, using the same weighted contrasts employed in the group analysis. We applied a more lenient statistical threshold (p < 0.05 uncorrected). Next, we determined the peak vertex for each subject’s four visual quadrant ROIs, i.e., the vertex with the highest t-value, for SBAV and CBA. To specifically assess the impact of macroanatomical alignment on the overlap of single-subject ROI peak vertices for each visual quadrant, we mapped all peak vertices per visual quadrant for SBAV and CBA. To quantify changes in the number of precisely overlapping single-subject peak vertices, we counted for each occipital vertex the number of peak vertices for SBAV and CBA. We performed this analysis in addition to the PM- and PDM-analysis to provide a more direct assessment and visualization of the effects of macroanatomical alignment on the spatial correspondence of single-subject ROIs.

## ACKNOWLEDGEMENTS

The authors are most grateful to Deliah Macht, Christina Raab and Leonie Winkler-Lauble for help with data acquisition. C.V Barnes-Scheufler was supported by a “main doctus” scholarship from The Polytechnic Foundation of Frankfurt am Main. E. Raspor was supported by a Research Grant for Doctoral Programmes in Germany from the German Academic Exchange Service (DAAD).

## AUTHOR CONTRIBUTIONS

All authors made substantial contributions to the conception or design of the work, or the acquisition, analysis, or interpretation of data. AR and RAB acquired funding. RAB, BP, MS and LR designed the experiment. MQ, CVB-S, LR and RAB acquired the data. ER and RG provided analytical tools. MQ, CS and RAB analyzed the data. MQ and RAB undertook the literature searches and wrote the first draft of the manuscript. All authors contributed to and revised the manuscript. All authors read and approved the final manuscript.

## DATA AVAILABILITY STATEMENT

The data that support the findings of this study are available from the corresponding author, R.A.B., upon reasonable request.

